# A shared limiting resource leads to competitive exclusion in a cross-feeding system

**DOI:** 10.1101/354282

**Authors:** Sarah P. Hammarlund, Jeremy M. Chacón, William R. Harcombe

**Affiliations:** Department of Ecology, Evolution, and Behavior, University of Minnesota, St. Paul, MN, USA; BioTechnology Institute, University of Minnesota, St. Paul, MN, USA

**Author notes:** both authors contributed equally (Sarah P. Hammarlund, Jeremy M. Chacón). Corresponding author: 140 Gortner Lab, 1479 Gortner Ave., St. Paul MN 55108, USA.

## Abstract

Species interactions and coexistence are often highly dependent upon environmental conditions. This is especially true for cross-feeding bacteria that rely on one another for essential nutrients. The addition of a cross-fed nutrient to the environment can release one species from its dependence on another, thereby altering the species’ interaction and potentially affecting coexistence. Using invasion-from-rare experiments with cross-feeding bacteria, genome-scale metabolic modeling, and classical ecological models, we explored the potential for coexistence when one cross-feeding mutualist becomes independent. We show that whether nutrient addition shifts an interaction from mutualism to commensalism or parasitism depends on whether the limiting nutrient can be metabolized by only one species or by both species. Furthermore, we show that coexistence is only lost when the interaction becomes parasitism, and the obligate species has a slower maximum growth rate. Surprisingly, models suggest that rates of cross-fed nutrient production have a negligible effect. These results contribute to an understanding of how resource changes, whether intentional or not, will manipulate interactions and coexistence in microbial communities.

## Introduction

Mutualistic interactions are abundant throughout the tree of life and across all ecosystems (Bronstein et al. 2004). The majority of flowering plants exchange pollen or nectar for pollination services, desert environments are dominated by legumes that house nitrogen-fixing rhizobia, and the fungal/algal symbioses that make up lichens cover up to six percent of Earth’s surface (Ollerton et al. 2011, Zahran 2001, Pennisi 2016). Mutualisms were historically viewed as being fixed—both species benefit from the other’s presence. However, most mutualisms do not fit this stereotype (Bronstein 1994, Sachs & Simms 2006, Hoeksema et al. 2010, Chamberlain et al. 2014). Mutualisms are often context-dependent, showing different magnitudes of mutual benefit or even qualitatively different interactions (e.g. from mutualism (+,+) to parasitism (+,−)) under different environmental conditions (Johnson & Graham 2013, Hoeksema & Bruna 2015). Because a change in an ecological interaction can lead to the loss of ecological stability, and potentially extinction, it is important to understand how context-dependence determines interactions and coexistence.

Context-dependence may be especially strong for mutualisms in which species exchange essential resources. When levels of resources in the environment change, the ‘mutualistic’ species may no longer depend on one another for these resources. Indeed, in plant-mycorrhizal fungi interactions and legume-rhizobia interactions, the obligacy and type of interaction can vary depending on levels of nitrogen or phosphorous in the environment (Heath & Tiffin 2007, Egerton-Warburton et al. 2007, Ehinger et al. 2009, Johnson 2010, Lau et al. 2012, Grman & Robinson 2013). If both species are abiotically provided with their resource requirements, the interaction shifts from mutualism (+,+) to either competition (−,−) or no interaction (0,0), the long-term outcomes of which are generally predictable, for example by R* (Tilman 1982). It is less clear, however, how a shift to unidirectional obligacy (when only one species is provided with resources) will change the qualitative interaction and potential for coexistence.

Mutualisms are common among microbes, who often reside in dense communities of many interacting species (Zelezniak et al. 2015). One example is bi-directional cross-feeding, where each species excretes a nutrient the other depends upon (Schink 2002, Pande & Kost 2017). Cross-feeding bacteria have been found in natural microbial communities and have evolved *de novo* in laboratory systems (Schink 2002, Goldford et al. 2017, Harcombe 2010, Hillesland & Stahl 2010). Metabolic dependencies may also explain why many species isolated from natural environments are not culturable (D’Souza et al. 2014). There is increasing interest in manipulating microbial communities by changing the availability of microbial nutrients (so-called “prebiotics;” Roberfroid et al. 2010). For such efforts to be predictable and successful, it is essential to understand how reduced obligacy of cross-feeding mutualisms changes coexistence of species.

In this work, we create two unidirectionally-obligate communities from a bacterial cross-feeding mutualism. We use experiments and modeling to determine how coexistence is influenced by resource availability, the physiological characteristics of the species (i.e. growth rate and production rate of cross-fed nutrients), and whether the system-wide limiting resource is private or communal. We test a hypothesis inspired by recent work (Hom & Murray 2014, LaSarre et al. 2016) that a community of two cross-feeding microbes will be stable if the faster growing species is limited by the slower growing species, but not vice-versa. We refer to this hypothesis as the “feed the faster grower” (FFG) hypothesis.

We examine the stability of unidirectionally-obligate communities using an experimental system of two human gut bacteria, *Escherichia coli* K12, and *Salmonella enterica* LT2 (Fig. 1A). Previously, this system was engineered and experimentally evolved into a metabolic mutualism in which our *S. enterica* strain secretes methionine in exchange for acetate byproducts from an auxotrophic *E. coli* (Harcombe 2010). In lactose minimal media the species form an obligate mutualism, but when methionine or acetate is added to the growth media, the species that requires that nutrient becomes independent of the other. In experiments and simulations, we find that the FFG hypothesis applies, but only under a restricted set of conditions. The slower growing species is lost upon feeding the faster grower only when the community is strongly limited by a communal nutrient that both species require. It is competition for this shared nutrient that causes exclusion of the slower species, regardless of the magnitude of any concurrent positive interactions.

**Figure 1:**
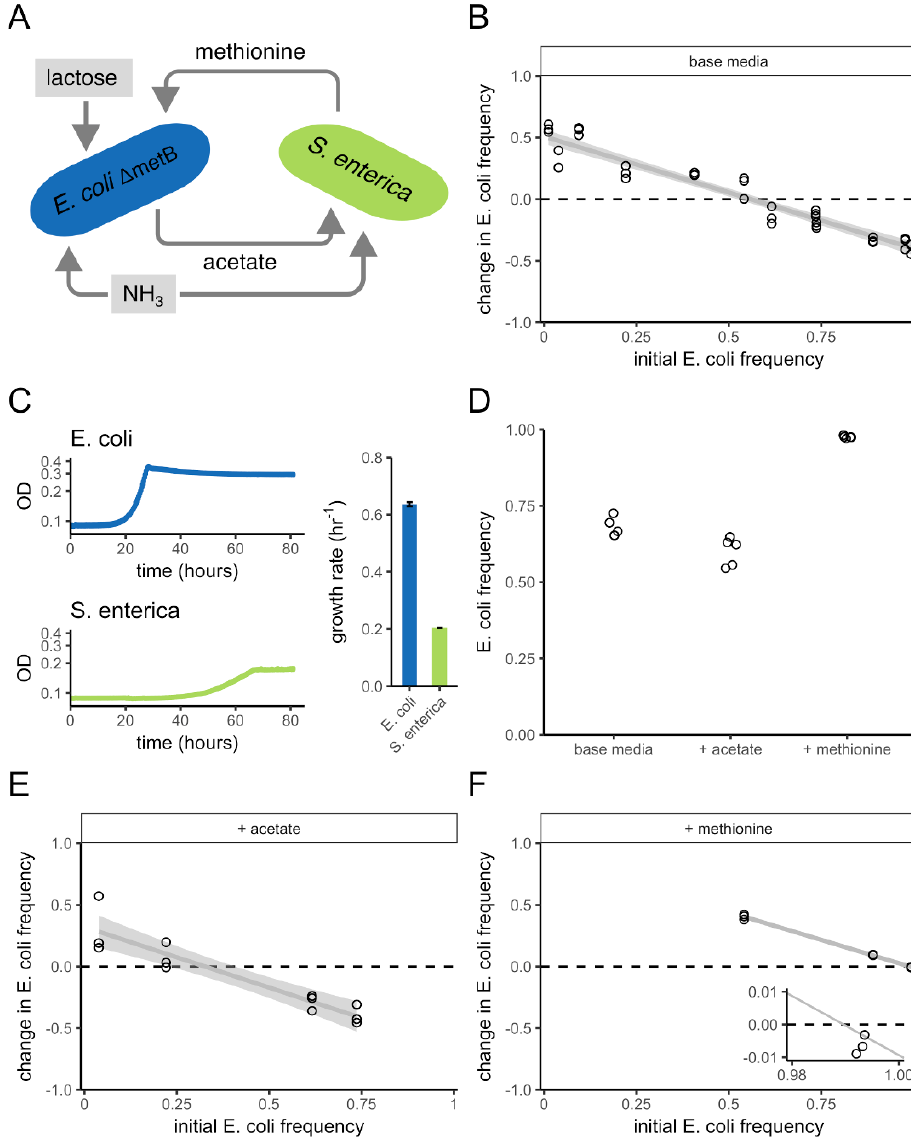
Nutrient additions to an obligate mutualism alter species ratios, but not coexistence. **(A)** Schematic showing the cross-feeding interaction between *E. coli* and *S. enterica*. *E. coli* consumes lactose, which *S. enterica* is unable to consume. *E. coli* excretes acetate as a by-product, which *S. enterica* consumes. *S. enterica* secretes methionine, required by *E. coli*. Both species consume ammonia. The resources provided in the base medium, lactose and ammonia, are highlighted in gray. **B)** The change in frequency of *E. coli* as a function of the initial *E. coli* frequency, in base media. Here, *E. coli* and *S. enterica* are obligate mutualists and can mutually invade from rare, as indicated by the downwards slope that crosses y = 0. Therefore, they can stably coexist (Chesson 2000). **C)** Growth curves showing optical density (absorption = 600nm) over four days of growth for *E. coli* (blue) and *S. enterica* (green) in monoculture in an environment where their required nutrient (methionine or acetate) is provided. *E. coli* has a faster growth rate than *S. enterica* (bar plot, error bars are standard deviations). N = five replicates per treatment. **D)** The final frequency of *E. coli* started from 0.5 in base media, or base media with acetate or methionine addition. The final frequencies upon either nutrient addition differed significantly from the final frequency in base media (see Results). **E)** The change in frequency of *E. coli* as a function of the initial *E. coli* frequency, in base media + acetate. **F)** The change in frequency of *E. coli* as a function of the initial *E. coli* frequency, in base media + methionine. When methionine is supplied, the initial *E. coli* frequency at which *E. coli* frequency decreases after four days of growth is much higher than in B or E, but coexistence is maintained, as shown in the inset plot where *E. coli*’s frequency decreases when started at an initial frequency of 0.997 (in the inset, points are jittered in the x-direction to show all replicates). In B, E, and F, gray lines are linear regressions and shaded gray regions are 95% confidence intervals.

## Results

### Reducing the dependence of the faster growing species on the slower growing species alters species frequencies, but not coexistence

A feature of obligate cross-feeding mutualisms is that the two species coexist at a stable frequency, indicated by both species’ ability to increase in frequency when initially rare (Shou et al. 2007). To confirm that *E. coli* and *S. enterica* co-cultures converge to a stable coexistence frequency when grown in lactose minimal media, we started co-cultures with initial *E. coli* frequencies ranging from 0.012 to 0.993, and found that both species, when initially rare, were able to increase in frequency after one four-day growth period (Fig. 1B).

We tested the impact of adding nutrients that released each species from reliance on the other. In our system, *E. coli* grows faster than *S. enterica* when grown in monoculture in environments that contain the metabolites used for growth in the mutualistic co-culture (Fig. 1C; one-tailed t-test, df = 4, p-value = 5.5e-9). Therefore, according to the FFG hypothesis, providing methionine in co-culture growth media, thereby releasing *E. coli* from its dependence on *S. enterica*, should destabilize the community, while adding acetate should not. When we added acetate to the lactose media and began co-cultures with equal population sizes of *E. coli* and *S. enterica*, we observed a slight decrease in *E. coli* frequency after four days of growth, compared to the lactose base media (Fig. 1D; two-tailed t-test, df = 4, p = 0.003). When we added methionine, we observed a large increase in *E. coli* frequency (two-tailed t-test, df = 4, p-value = 1.8e-5), indicating that the FFG hypothesis may apply.

However, when we conducted invasion-from-rare experiments to assess coexistence, coexistence was maintained in both the acetate addition (Fig. 1E) and the methionine addition (Fig. 1F) environments. Under methionine addition, *E. coli* dominated the community, but the new species ratio was stable, i.e. *S. enterica* was still able to invade from rare (Fig. 1F inset; one-tailed t-test, df = 2, p-value = 0.02). This result suggests that feeding the faster grower alters the mutualism to the extent that species ratios change, but does not by itself destroy coexistence.

### Exposure of underlying resource competition by limiting a shared nutrient causes the faster growing species to exclude the slower growing species

Classically, two species compete to the point of competitive exclusion if a single communal resource that both species require is limiting (Gause 1934, Hardin 1960, Hutchinson 1961). Therefore, we hypothesized that reducing the concentration of a communal nutrient until it is limiting, in addition to feeding methionine to *E. coli*, could cause exclusion of *S. enterica*. In our system, the nitrogen source ammonia is a communal resource that both species require, so we examined the effect of limiting ammonia on coexistence when either species was provided with its cross-fed nutrient.

We first generated predictions using dynamic multi-species genome-scale metabolic modeling. Metabolic modeling allows us to set the abundance of every metabolite in the simulated environment, and growth of the species emerges as a function of this environment and their metabolic networks. Using well-curated models of our experimental system (Harcombe et al. 2014, and see Experimental

Procedures), we first mimicked the environment in our lab experiments. The species coexisted in a stable mutualism in an environment initiated without acetate or methionine (Fig. 2A). Like in our experiments, both species could invade from rare when acetate (Fig. 2B) or methionine (Fig. 2C) was supplied. We then repeated these simulations with limiting ammonia concentrations (1/20^th^ of the initial concentration used in experiments), which we hypothesized would change the interaction to parasitism. This change switched the limiting nutrient from lactose to ammonia and reduced productivity of the community (data not shown). We still saw coexistence in the base media (Fig. 2D) and when acetate was supplied (Fig. 2E), but *E. coli* excluded *S. enterica* when methionine was supplied (Fig. 2F), indicating that coexistence is lost when slower-growing *S. enterica* both depends upon *E. coli* and competes with it for a limited resource, simultaneously.

**Figure 2:**
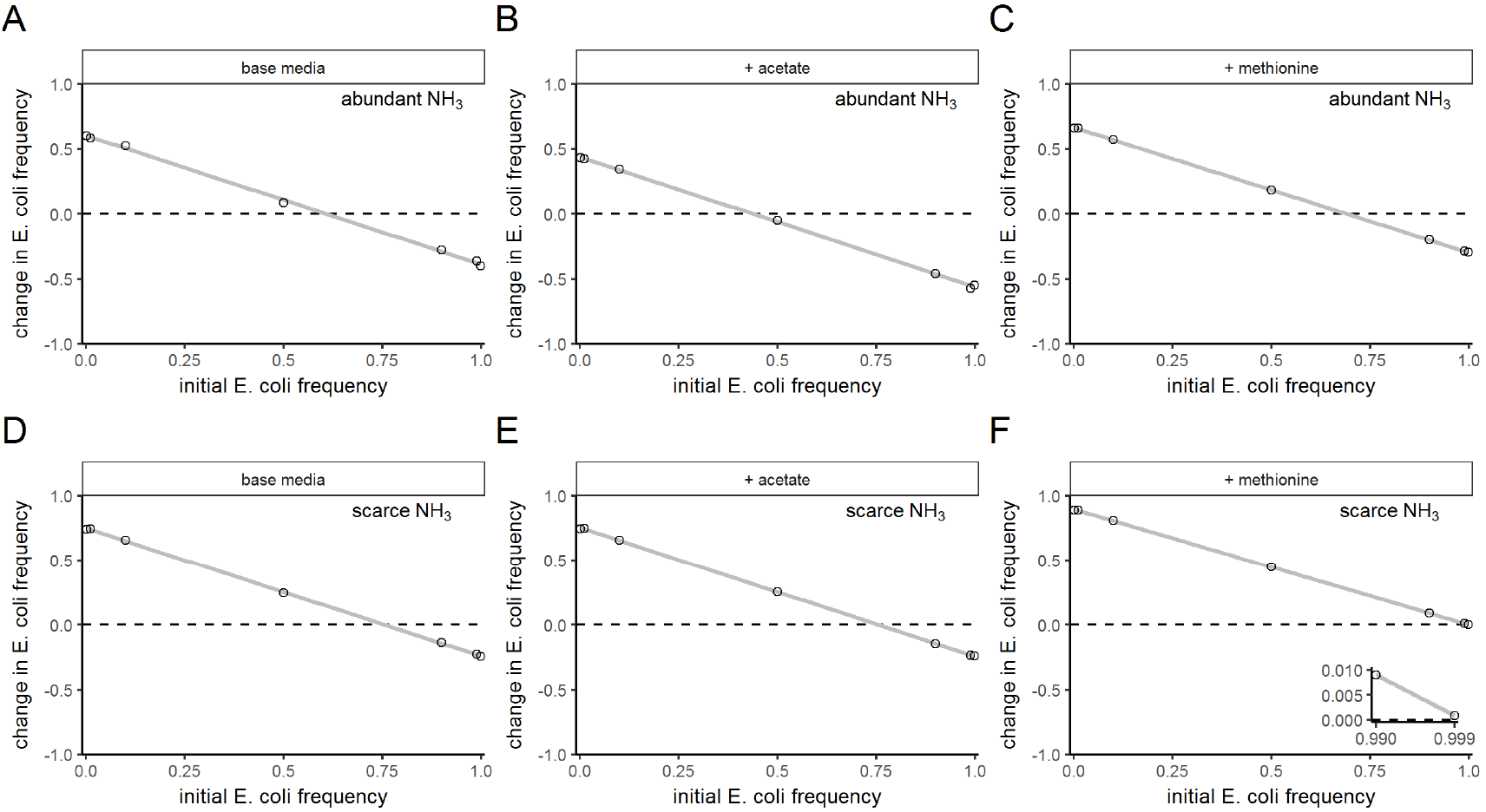
Dynamic metabolic modeling predicts that coexistence of an obligate cross-feeder with its nutrient supplier depends on species identity and the abundance of the communal nutrient, ammonia. Invasion-from-rare experiments using dynamic metabolic models in environments with non-limiting **(A-C)** or strongly limiting **(D-F)** concentrations of ammonia. Environments in **(A, D)** contained base media, **(B, E)** contained acetate added to base media, and **(C, F)** contained methionine added to base media. The y-axes show the change in frequency *E. coli*, as a function of the initial *E. coli* frequency. As in Fig. 1, trends that cross y = 0 with a downwards slope show mutual invasibility and therefore coexistence. The inset in **(F)** shows that *S. enterica* could not invade even when simulations began with a very high frequency of *E. coli*, because the *E. coli* frequency still increased. Lines are least-squares linear fits.

Because the hypothesis was supported by our mathematical predictions, we tested the effect of limiting ammonia in our experimental system. Ammonia limitation by itself did not affect coexistence when each species relied on metabolites from the other species (Fig. 3A). Adding acetate to low-ammonia media also did not affect coexistence (Fig. 3B; one-tailed t-test, df = 2, p-value = 0.047). However, when we added methionine to low-ammonia media, *E. coli* increased in frequency from any starting frequency, showing that *S. enterica* is unable to survive (Fig. 3C; one-tailed t-test, df = 2, p-value = 4.0e-5; see inset).

### A simple ecological model shows that competitive exclusion is driven by growth rate differences and strong competition for a communal limiting resource

Next, we explored the generality of these results and controlled for a number of correlated variables which were difficult to study experimentally. For example, in our system, *E. coli* is the faster grower, but it is also the ‘less generous’ metabolic partner, providing fewer cell-equivalents of acetate per unit of growth than *S. enterica* provides of methionine (as evidenced by the community being biased towards *E. coli* when grown in lactose base media). Second, in a sense, *E. coli* ‘governs’ the community, because it performs the first step of metabolism of the energy resource, lactose.

**Figure 3:**
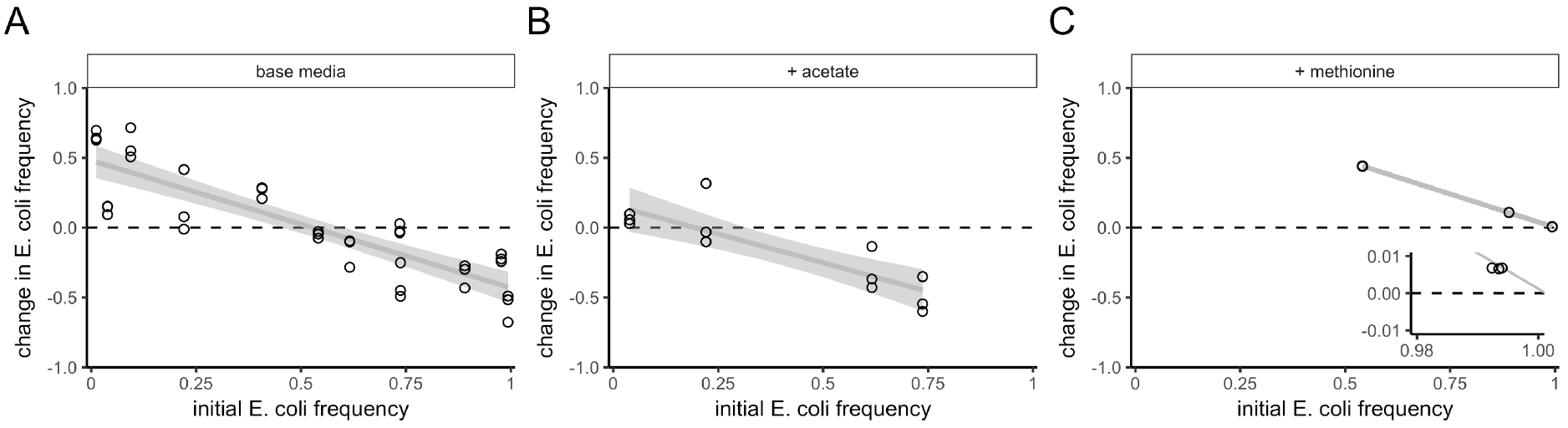
In *in vitro* experiments, ammonia limitation causes competitive exclusion of *S. enterica* when *E. coli* is made independent by addition of methionine. All panels show invasion-from-rare experiments, *in vitro*, in environments with limited ammonia and acetate or methionine addition. The y-axes show the change in frequency of *E. coli*, as a function of the initial *E. coli* frequency. As in Fig. 1, trends that cross y = 0 with a downwards slope show mutual invasibility and therefore coexistence. The environment in **(A)** contained the base media, **(B)** contained acetate added to base media, and **(C)** contained methionine added to the base media. The inset in **(C)** shows that *S. enterica* could not invade under ammonia limitation and methionine supplementation, which is consistent with the dynamic genome-scale metabolic modeling. The lines are least-squares linear fits, with the shaded region enclosing 95% confidence intervals.

To disentangle these factors and explore the effects of acetate and methionine addition, nitrogen limitation, and relative growth rate differences, we turned to a theoretical exploration using a system of differential equations.

The initial system of equations contained twelve parameters: a maximum growth rate for each species, a Monod constant and an efficiency term for each resource, and production terms for methionine by *S. enterica* and acetate by *E. coli*. Because we were primarily interested in the initial resource values, relative growth rates, and relative rates of nutrient production, we scaled model parameters into the following equations (see Supplemental Methods for scaling procedure):

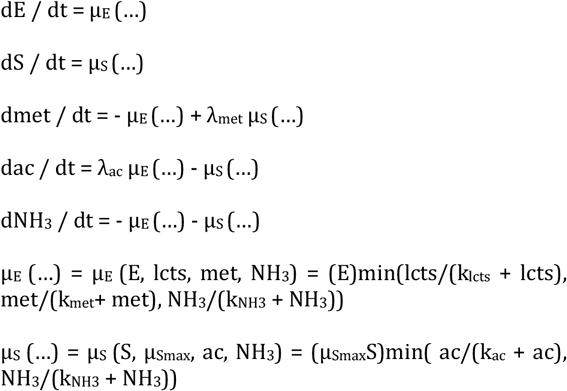

Here, *E* and *S* are the cell counts of each species. Resources (lcts = lactose, ac = acetate, met = methionine, and NH_3_ = ammonia) are in units of ‘cell-equivalents’ meaning one resource of each type required is used to create one cell. Monod constants (k) are also in units of cells. *E*’s maximum growth rate was removed by scaling time, and *S*’s maximum growth rate (μ_Smax_) is proportional to *E*’s. Production terms (λs) are in units of cells/cell, i.e. are unit-less, and can be considered to be the number of cells of the other species that can be supported by growth of one cell of the focal species.

Growth is governed by Monod saturation rates, and a Liebig’s law of the minimum is applied so that only one resource is ever limiting for a species at a time. This scaling allowed us to clearly focus on the initial resource abundances provided, the relative growth rates, and the rates of production of the cross-fed nutrients.

We began with a situation that approximates our experimental system to ensure the model is capable of reproducing our empirical findings. *S* has a maximum growth rate of 0.5 that of *E*, and *E* produces fewer cell-units of acetate than *S* produces of methionine (based on final yields in mutualistic co-cultures), so we set λ_ac_ = 0.4 and λ_met_= 7.3. With these parameters, we were able to recapitulate the experimental and genome-scale simulation results (Fig. S1).

Satisfied that the model can recapitulate our experimental and genome-scale data, we then explored a wider parameter space. To disentangle the correlated difference in cross-fed nutrient production with the difference in growth rates, we set both production terms, λ_ac_ and λ_met_, to 1.5. The product of the production terms must be greater than one to sustain the mutualism (Shou et al. 2007, Fig. S2). We kept *S*’s growth rate at half of that of *E*, to be able to test the FFG hypothesis. To understand how different degrees of mutual obligacy affected coexistence, we swept through many initial values of met, which reduces *E*’s dependence on *S*. To learn how this interacts with co-limitation by NH_3_, we also swept through different initial values of NH_3_. To assess coexistence, we started simulations with *S* rare (1%) and tracked whether *S* could invade. Finally, we also ran monoculture simulations of *E*, to compare *E*’s final density in the presence vs. absence of *S* and find the qualitative ecological effect (+,0, or −) *S* had on *E*.

**Figure 4:**
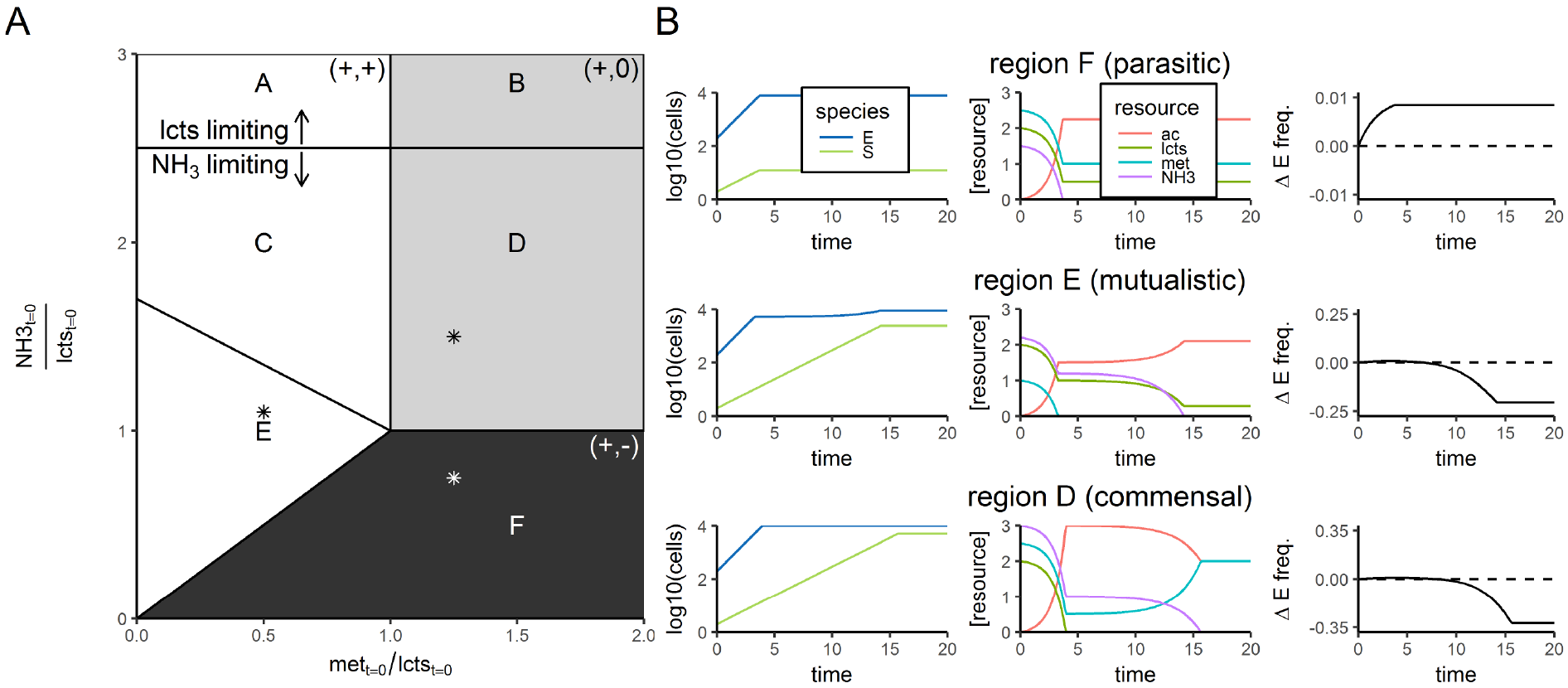
Simulated effect of initial methionine and NH_3_ concentrations on the ability of a slower-growing obligate cross-feeding species to coexist with its nutrient supplier. **A)** Diagram showing whether *S* could increase in frequency when rare, and the net ecological interaction between *E* and *S*, for batch culture simulations initiated at many different values of methionine and NH_3_. In these simulations, μ_Smax_ = 0.5, lcts_t=0_ = 10000 cell equivalents, λ_met_ = λ_ac_ = 1.5, all k_m_ = 1e-3 cell equivalents, *E*_t=0_ = 198, and *S*_t=0_ = 2. Different polygons enclose qualitatively different results in terms of what nutrients ran out (see Table 1). Colors indicate the net ecological interaction. The first symbol indicates the net effect of *E* on *S*, the second symbol the net effect of *S* on *E:* white = mutualistic interaction (+/+); gray = commensal interaction (+/0); black = parasitic interaction (+/−). In addition, in the black region, *S* could not increase from rare, but it could in the white and gray regions. The data used to make this diagram come from 400 simulations, at evenly-spaced x-and-y initial conditions. **B)** Representative time series from three regions where NH_3_ limits the system, with initial values taken at the locations marked by asterisks. Resource concentrations, in units of cell equivalents, are 5000 times the y-axis value. The third column shows the change in *E* frequency, from an initial frequency of 0.99. A negative change indicates that *S* can invade from rare, meaning both species can coexist.

Six qualitatively different results occur when different amounts of nutrients are provided, which are shown in Fig. 4 and summarized in Table 1. Briefly, coexistence is prevented when the net interaction is parasitic (+,−), which occurs when the amount of NH_3_ provided is less than the amount of growth that *E* could sustain on the environmentally-provided lcts and met (region F). Coexistence is possible in all other regions, regardless of whether the net interaction is mutualistic or commensal, or whether productivity is low and there is competition for NH_3_.

**Table 1:** Coexistence potential and interaction type in different nutrient environments. The regions (column 1) correspond to those depicted in Fig. 4A. Coexistence was measured by invasion-from-rare experiments. The net effect on *E* was measured by comparison with monoculture. Resource exhaustion was measured when dynamics ceased.

| Region | Coexistence | Net interaction<br>(effect of $E$ on $S$, effect of $S$ on $E$) | Resources<br>that run out | $E$ limited by | $S$ limited by | Why the net<br>effect on $E$? | Does<br>competition<br>for $\text{NH}_3$ affect<br>yields? |
| --- | --- | --- | --- | --- | --- | --- | --- |
| A | Yes | (+,+) | lcts | lcts | ac | $E$ gets met from $S$ | No |
| B | Yes | (+,0) | lcts | lcts | ac | Environment<br>provides<br>sufficient met | No |
| C | Yes | (+,+) | $\text{NH}_3$, lcts | met, then<br>lcts | $\text{NH}_3$ | $E$ gets met from $S$ | Yes |
| D | Yes | (+,0) | $\text{NH}_3$, lcts | lcts | $\text{NH}_3$ | Environment<br>provides<br>sufficient met | Yes |
| E | Yes | (+,+) | $\text{NH}_3$, met | met, then<br>$\text{NH}_3$ | $\text{NH}_3$ | $E$ gets met from $S$ | Yes |
| F | No | (+,-) | $\text{NH}_3$ | $\text{NH}_3$ | $\text{NH}_3$ | Competition for<br>$\text{NH}_3$ | Yes |

If we repeat the initial resource value sweep but instead of providing met for *E*, we provide acetate for *S*, we see a very different picture: *E* can always invade from rare, showing that *E* and *S* coexist, regardless of NH_3_ levels (Fig. S3). In other words, when a faster-growing species obligately depends on a slower-growing species, the obligate species can always invade. There is still a region where the relationship is parasitic: where there is sufficient ac to sustain *S*, yet *E* competes with *S* over NH_3_ (Fig. S3B, red region). Interestingly, despite NH_3_ being the communal limiting resource in this case, the species coexist. This is because *E*’s higher maximum potential growth rate, coupled with its dependence on *S*, allow *E* to keep pace with *S*.

These results suggest that coexistence of an obligate species with its nutrient provider depends on both relative growth rates and the degree to which productivity is limited by the concentration of a communal nutrient. Next, we tested the effect of relative growth rate more directly by running simulations in environments that cause a parasitic interaction (region F in Fig. 4), and sweeping across relative growth rates. We also alter which species is obligately dependent upon the other in order to disentangle the effect of *E* being the lactose metabolizer. We find that there is a hard switch from exclusion to coexistence that occurs when the growth rate of the obligate species is greater than or equal to the growth rate of the independent species (Fig. 5A). This effect is independent of whether *E* or *S* is the obligate cross-feeder.

**Figure 5:**
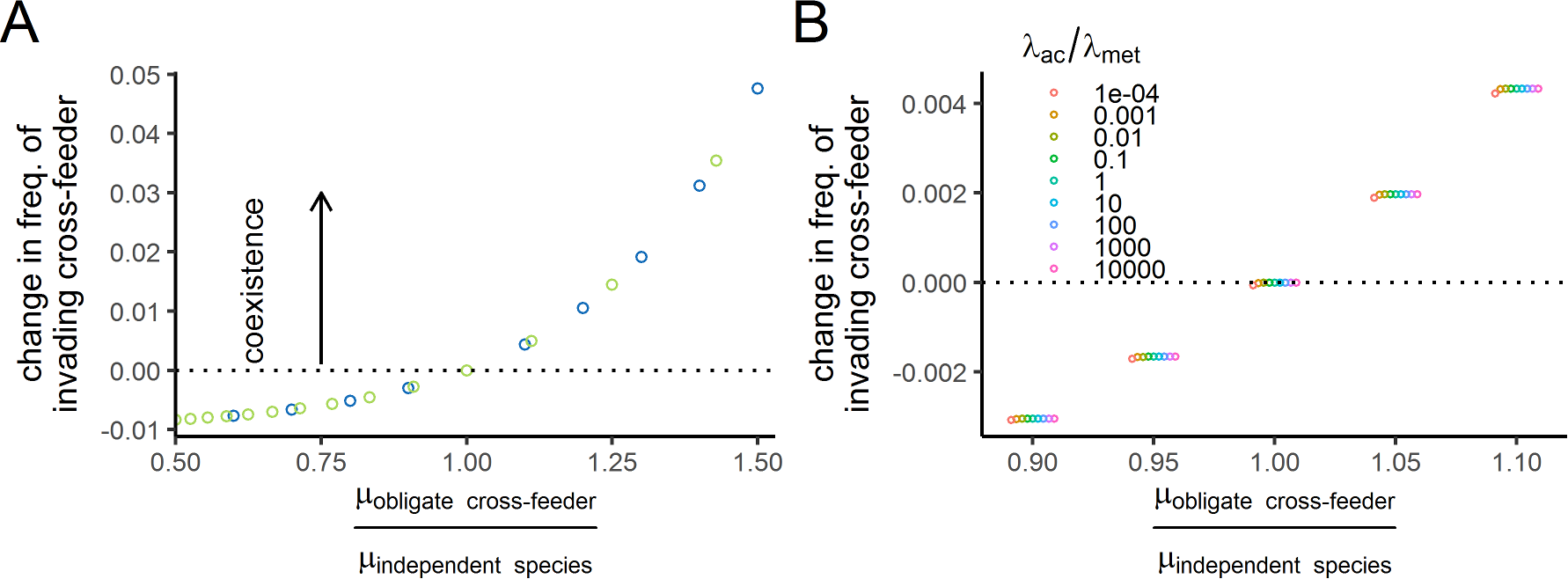
Simulations predict that relative growth rates, but not cross-fed nutrient production rates, determine whether an obligate cross-feeder can coexist with an independent nutrient supplier. These panels show the results of invasion-from-rare simulations when the net ecological interaction is parasitic. Therefore, all simulations were run under severe NH_3_ limitation and with excess cross-fed nutrient provided to one species. **A)** The change in frequency of an invading cross-feeder plotted over the cross-feeder’s relative growth rate. Points above the dotted line indicate successful invasion and therefore coexistence of the cross-feeder with the nutrient provider. The color of the circles indicates which species is the obligate cross-feeder (blue for *E*, green for *S*). When *E* was the cross-feeder, *S* was provided excess ac, and when *S* was the cross-feeder, *E* was provided excess met. **B)** The change in frequency of an invading cross-feeder as a function of both relative growth rate (x-axis) and the relative "generosity" of the species (relative production terms, λs, colored points). In these simulations, the obligate cross-feeder, *S*, depends on ac from *E*, which produces it at a rate of λ_ac_. The product of the λ values was held constant at 2.25.

Finally, we tested the effects of the “generosity” of the two species: specifically, how much resource they provide to the other species (λ_ac_ and λ_met_). In our experiments, λ_ac_ is lower than λ_met_. Does varying this ratio alter coexistence? Surprisingly, even extreme values of this ratio have a negligible effect (Fig. 5B), although it is possible that an effect would emerge if the species were poorer at taking up resource at low concentrations (i.e. had higher km values). In these simulations, where both species were efficient at taking up all nutrients, only extremely selfish *E* (low λ_ac_/ λ_met_) impairs invasion by *S*, and only when their growth rates are equal.

## Discussion

We explored how changes in the resources in an environment altered the interaction between two cross-feeding bacteria, and the consequences of that altered interaction for coexistence. We started with an obligate cross-feeding mutualism between *E. coli* and *S. enterica*. Then, using experiments and models, we made one species metabolically independent, by adding either methionine or acetate, and tested whether both species could continue to coexist. We found that acetate addition, which made *S. enterica* independent, always led to coexistence. Methionine addition, which made *E. coli* independent, sometimes led to competitive exclusion of *S. enterica*. *S. enterica* exclusion depended upon it having a slower growth rate than *E. coli, and* on the productivity of the system being set by ammonia rather than lactose. This was because sufficiently strong ammonia limitation caused the interaction to shift to parasitism when methionine was added, whereas lactose limitation caused the interaction to shift to commensalism. Interestingly, the relative production of cross-fed nutrients had little impact on the prevalence of coexistence. Below, we discuss our two key conclusions: First, microbial interactions can readily be shifted from mutualism to commensalism or parasitism by altering nutrient conditions. Second, whether release from mutual dependency leads to coexistence or competitive exclusion is highly context-dependent, depending on both on the type of interaction and relative growth rates.

Whether a cross-feeding mutualism shifts to commensalism or parasitism upon release of a dependency is determined by the nature and relative concentration of the resource that limits system productivity. In our system, productivity was limited either by lactose, which was private to *E. coli*, or by ammonia, which was communal. The sign and strength of the mutualism (without addition of a cross-fed nutrient) were the same regardless of which metabolite limited system productivity. However, the impact of adding cross-fed metabolites had categorically different ecological effects under the two limiting resources. Strong ammonia limitation led to parasitism when cross-feeding became unidirectional because any growth of the dependent species came at the cost of the independent species’ yield. This reduction in final yield was eliminated when system productivity was set by the private resource, lactose. Hoek et al. (2016) also found that the ecological interaction between cross-feeding strains could readily shift upon addition of cross-fed nutrients. In that study, addition of a high concentration of cross-fed nutrients always led to negative interactions. This finding is consistent with our results, because their cross-feeding organisms were two nearly isogenic strains of *Sacchromyces cerevisiae*, and therefore every resource except the cross-fed nutrients was communal. In general, as Hoek et al. also suggest, the more similar the metabolism is between cross-feeding strains, the more likely it may be that nutrient addition will shift a mutualism to a negative interaction. Consistent with this idea, mathematical predictions suggest that cross-feeders must each specialize to coexist, because otherwise competition will cause the system to collapse (Doebeli 2002, Pfeiffer & Bonhoeffer 2004, Gudelj & Rosenzweig 2016, McCully et al. 2017). It will be interesting to examine whether stable cross-feeding interactions in natural systems are more common between species with smaller degrees of niche overlap.

Whether an obligate cross-feeder can coexist when acting as a parasite is determined by growth rate. When ammonia was sufficiently limiting, our system could be switched to parasitism by adding either acetate or methionine. However, the two additions had distinct outcomes on coexistence: acetate addition allowed coexistence, but methionine addition did not. This divergence in outcomes was determined by the relative maximum growth rate of the species released from metabolic constraint. If the slower growing species (*S. enterica*) was released, coexistence was maintained because the better competitor for ammonia (*E. coli*) could not outgrow the species from which it cross-fed. However, when the faster species was released from dependency, the slower grower was outcompeted. This finding is consistent with the feed the faster grower hypothesis suggested by Hom and Murray (2010) and LaSarre et al. (2017), though it is important to reiterate that feeding the faster grower did not result in destabilization under our commensalism regime. Furthermore, the destabilization seen in these previous studies was driven by a different underlying mechanism. In those studies, when the faster-growing species was supplied its cross-fed nutrient in the growth media, it produced toxic organic acids that harmed the slower-growing species and allowed the faster grower to overtake the community (shown in LaSarre et al. 2017, and likely occurring in Hom and Murray 2010). This work suggests that changes in nutrients are likely to have much more dramatic effects on community diversity when species differ in their maximum potential growth rates.

Our results highlight the importance of considering traits (a mechanistic approach) rather than simply net ecological interactions (a phenomenological approach) to understand community dynamics. Net ecological interactions are often measured by comparing growth of a focal species in the presence vs. absence of a second species (e.g. Vetsigian et al. 2011, Foster & Bell 2012). In our system, when methionine was added and ammonia was sufficiently limiting, *S. enterica* gained a net positive effect from *E. coli* due to acetate cross-feeding. Despite the benefit gained from *E. coli*’s presence, *S. enterica* was competitively excluded from the community. This occurred because competition for ammonia was an underlying negative component of the net positive interaction, and this underlying component drove the system. Therefore, predicting coexistence of *E. coli* and *S. enterica* required a detailed understanding of the species’ traits (i.e. the species’ growth rates and which nutrients they consume). This result would have likely been missed in a coarse-grained Lotka-Volterra modeling approach (as Momeni et al. 2017 suggest). In contrast, by using a trait-based approach that explicitly incorporates nutrient consumption and growth rates, our metabolic and ecological modeling approaches were able to accurately predict coexistence outcomes. Measuring net ecological interactions is simpler than determining which traits are relevant and measuring them. However, improvements in our ability to predict species’ traits from their genomes should reduce this workload (e.g. Machado et al. 2018 for primary metabolic traits, Weber et al. 2015 for secondary metabolic traits). Furthermore, a trait-based approach allows for prediction across many different environmental conditions. Within specific nutrient environments, phenomenological approaches have been successful predictors of coexistence (Friedman et al. 2017, Venturelli et al. 2018). However, because different environments produce different net ecological interactions, but do not affect species’ intrinsic traits, we argue that trait-based approaches will be an integral tool for predicting microbial community assembly and dynamics across a range of environments.

Finally, our results can inform efforts to manipulate microbial communities. Understanding how communities respond to changing environments is important for managing microbial ecosystems with prebiotics, as prebiotic nutrient addition is unlikely to be sustained indefinitely (Rastall & Gibson 2015). If a beneficial bacterium typically in an obligate cross-feeding mutualism is fed a prebiotic, and its partner is lost, cessation of prebiotic addition will result in the loss of the beneficial species. More broadly, our results demonstrate that a detailed understanding of who cross-feeds what from whom will significantly enhance our ability to manipulate microbiomes using prebiotics, with fewer off-target effects.

## Experimental Procedures

### Strains and media

We used *Escherichia coli* strain K12 with a Δ*metB* mutation conferring methionine auxotrophy and a *Salmonella enterica* serovar Typhimurium LT2 strain containing mutations in *metA* and *metJ* that cause over-production of methionine (Harcombe 2010; Douglas et al. 2016, 2017). In lactose minimal medium, *E. coli* consumes lactose and excretes acetate as a byproduct. *S. enterica* consumes this acetate and secretes methionine. For all experiments, strains were grown in a modified Hypho minimal medium base with lactose as the primary carbon source, containing 2.78mM lactose, 14.5mM K_2_HPO_4_, 16.3mM NaH_2_PO_4_, 0.814mM MgSO_4_, 3.78mM Na_2_SO_4_, trace metals (1.2uM ZnSO_4_, 1uM MnCl_2_, 18uM FeSO_4_, 2uM (NH_4_)_6_Mo_7_O_24_, 1uM CuSO_4_, 2mM CoCl_2_, 0.33um Na_2_WO_4_, 20uM CaCl_2_) and either 3.78mM (non-limiting) or 0.378mM (limiting) [NH_4_]_2_SO_4_. Where indicated, 80μM of L-methionine, or 12mM of acetate was added. Before beginning experiments, the strains were streaked from freezer stocks onto Nutrient Broth plates and grown for two days. Single colonies were then picked into 10mL of Hypho minimal media with methionine supplied for *E. coli* and 5.56mM glucose instead of lactose for *S. enterica*, in flasks, which were grown for two days, shaking at 30°C. Around six hours before beginning experiments, cultures were diluted 10-fold, and once cloudy, diluted into a carbon-free Hypho buffer based on OD600, to equalize cell densities. Experimental cultures were inoculated 2 × 10^4^ cells for monocultures or 4 × 10^4^cells for cocultures, with the desired frequency of *E. coli* relative to *S. enterica*.

### Growth assays

To measure growth rates in monoculture, methionine was supplied for five replicate *E. coli* monocultures and acetate was supplied for five replicate *S. enterica* monocultures. These monocultures were grown in 96-well plates in a Tecan InfinitePro 200 at 30°C, and measurements of OD600 were taken every 20 minutes, shaking at 432 rpm between readings. To obtain growth rate estimates, we fit growth curves to a Baranyi function (Baranyi & Roberts 1994) by obtaining nonlinear least-square estimates and using the growth rate parameter estimate.

In co-culture experiments, cultures were grown in Hypho media with either no addition (base media), acetate addition, or methionine addition, and with non-limiting or limiting ammonia concentrations, in 96-well plates at 30°C in a Bellco mini-orbital shaker, shaking at 450rpm. After four days of growth, the final yield of each species was measured through serial dilution and plating onto species-specific Hypho media with 1% agar. For *E. coli*, media contained methionine, and for *S. enterica*, lactose was replaced with 16.6mM glucose. X-gal (0.05% v/v) was added to plates to further help distinguish *E. coli* colonies (blue) from *S. enterica* colonies (white). All co-culture experiments had three or five replicate cultures for each initial frequency.

### Invasion-from-rare assays

We used mutual invasibility as our criterion for whether the two species could coexist—coexistence is possible when both species are able to increase in frequency from rare (Chesson 2000, Wright & Vetsigian 2016, Friedman et al. 2017). Cells in log phase (see above) were diluted into wells of a 96-well plate at different frequencies. The lowest initial frequency of *E. coli* was 0.012, and the highest was 0.997. After 4 days of growth, the yield of each species was measured by plating, as described above.

### Genome-scale metabolic modeling

The dynamic, multi-species genome-scale metabolic modeling was carried out in the COMETS platform (Harcombe et al. 2014). This simulates growth of a community by iterating over small time-steps. An environment, which is the abundance and identity of the nutrients the species’ models can use or produce, is pre-defined at the onset of the simulation. In a time-step, flux-balance analysis is used to determine the maximum amount of growth each species is capable of, as well as the nutrients used and produced to obtain this growth. The growth is constrained by the ability of the metabolic model to flow available external resources to the biomass objective reaction, as well as by the maximum uptake possible by that species, which is determined by Michaelis-Menten kinetics (vmax = 10, km = 5e-6 for all simulations). Once the flux-balance-analysis is solved in a time-step, the biomass growth and nutrient changes are scaled by the current amount of biomass and by the duration of the time-step, then the environment is adjusted by these changes. The species update sequentially each time step, and the order of species is randomized each time step. See Harcombe et al. (2014) for additional details.

The genome-scale models are previously published modifications of the standard *E. coli* and *S. enterica* metabolic models (Harcombe et al. 2014). The *E. coli* model, ijo1366 (Orth et al 2011), was modified by setting the bounds on the cystathionine gamma-synthase reaction (a reaction in the methionine biosynthetic pathway) to zero, which causes this model to require methionine from the external environment to be able to grow. The *S. enterica* model, (iRR_1083, Raghunathan et al. 2009), was modified so that for every unit of growth a proportional amount of methionine was excreted into the environment. These models can grow together in a COMETS simulation with lactose, ammonia, oxygen and trace minerals present, but neither can grow alone.

In all simulations, the following compounds were available in unlimited concentrations: Ca2[e], Cobalt2[e], cl[e], cu[e], fe2[e], fe3[e], k[e], mg2[e], mn2[e], mobd[e], ni2[e], o2[e], pi[e], so4[e], and zn2[e] (following BIGG notation). Lactose was supplied at the same concentration as in experiments, 2.78mM (equivalent to 1g/L). NH_4_, methionine, and acetate were also supplied at the same concentrations as experiments, with abundant NH_4_ = 7.56mM, scarce NH_4_ = 0.378mM, methionine = 80μMwhen in excess, and acetate = 12mM when in excess. In COMETS, molarity is determined by setting the mmols of each resource, and the “spacewidth,” which is the length of a three-dimensional box that determines the experimental volume. To mimic a 200μl well, spacewidth = 0.5848035cm. Time step = 0.01 hours. Biomass in flux-balance analysis is in units of grams dry weight. Initial total biomass = 5.6e-8g (roughly 10^4 cells), and divided into *E. coli* and *S. enterica* depending on the frequency tested in the given experiment.

### Differential equation ecological modeling

The scaled ecological model is shown in the Results, and all relevant parameters / initial values are stated either in results or figure legends. The ODE system was solved using the deSolve package in R, which used the lsoda solver to numerically integrate. All simulations were solved for sufficient duration to ensure dynamics had ceased. A maximum time step was set at 0.1 time units. During integration, relative tolerance (rtol) was set = 1 × 10^−10^ and maxsteps = 1 × 10^6^.

Because the resources were abiotic and the system was closed, the resource equations were recast as algebraic equations which improved the stability of the numerical solver. For example, lactose became a function of initial amount and the amount of growth by E: Lactose(time = t) = lactose(time = 0) – (E(time = t) – E(time = 0)). Occasionally, due to the discrete nature of numerical integration, a resource concentration would drop to slightly below zero. These events were tested for and if they occurred, the concentration was set = 0. Such small numerical issues never affected growth more than 1/100 of a cell’s worth of growth.

### Statistics

All analysis was done in R.

## Acknowledgements

The authors thank B. Adamowicz, L. Fazzino, J. Anisman, A. Behling, F. Isbell, C. Daws, and the UMN theory group for useful discussions. S. Hammarlund was funded by a NSF Graduate Research Fellowship and J. Chacón through NIH (GM121498-01A1).

## Supplemental Figures

**Figure S1.**
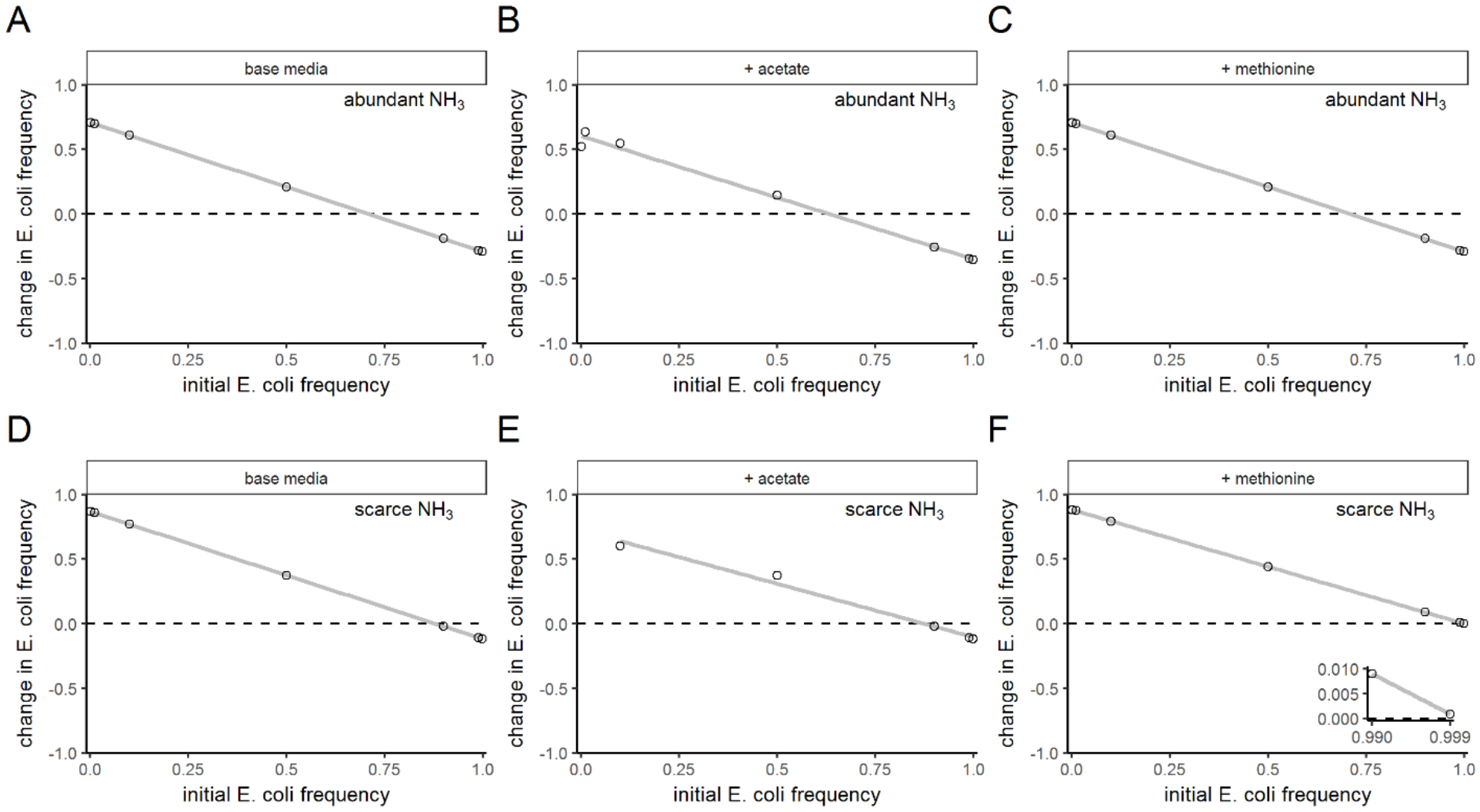
The scaled ecological model can recapitulate the coexistence patterns observed in experiments and genome-scale metabolic models. Invasion-from-rare simulations were conducted using the scaled model with parameters similar to our experimental system, in which *E* is the less generous partner (λ_ac_ = 0.4) compared to *S* (λ_met_ = 7.3) and *S* grows at approximately half the maximum growth rate of *E* (μ_Smax_ = 0.5). All panels show the change in frequency of *E* over the initial *E* frequency. lcts_t=0_ = 10000 for all simulations. NH_3t=0_ = 15520 (non-limiting) in **(A-C)** but is limiting (NH_3t=0_ = 705) in **(D-F)**. In **(A,D)**, met_t=0_ = ac_t=0_ = 1, to kick-start the mutualism. In **(B,E)** met_t=0_ = 1 and ac_t=0_ = 4110 (making *S* independent). In **(C,F)** met_t=0_ = 10000 and ac_t=0_ = 1 (making *E* independent). The inset in **(F)** shows that *E* increases in frequency even when abundant, meaning *S* is not able to coexist with *E*.

**Figure S2.**
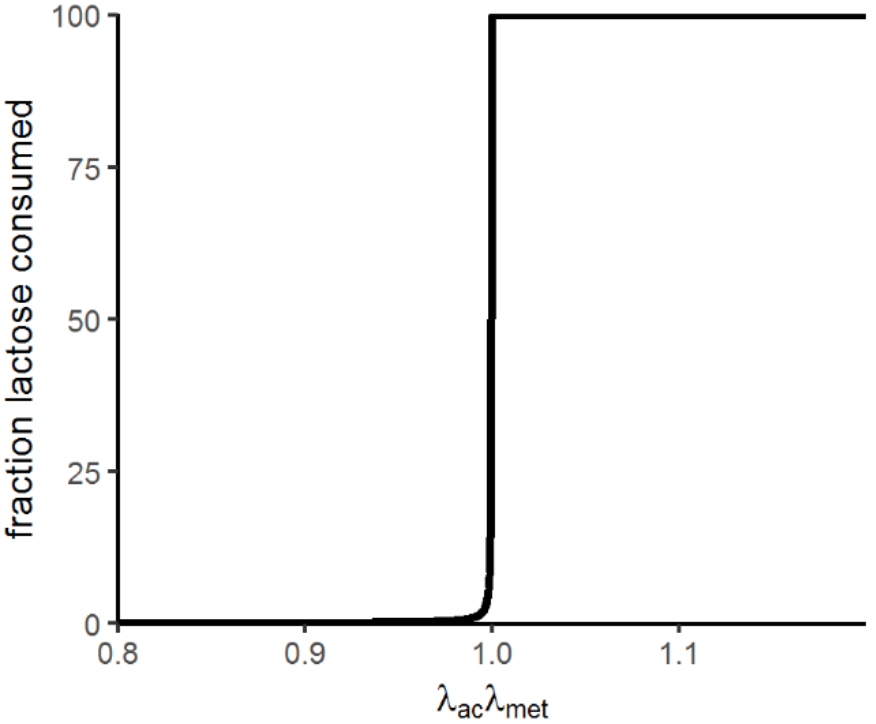
The product of the cross-fed nutrient production parameters must by greater than one to sustain mutualism. Results of simulations with excess NH_3_ and 10 units of acetate initially present to “jumpstart” the mutualism. The x-axis shows the product of the two production rate parameters. The y-axis shows the percent of the lactose that was consumed. When this y value is < 100%, production of cross-fed nutrients is too low to sustain growth, and met and acetate run out. When this value is > 100%, production of cross-fed nutrients sustains growth and lactose limits the system. In these simulations, μ_E,max_ = 2 μ_S,max_.

**Figure S3.**
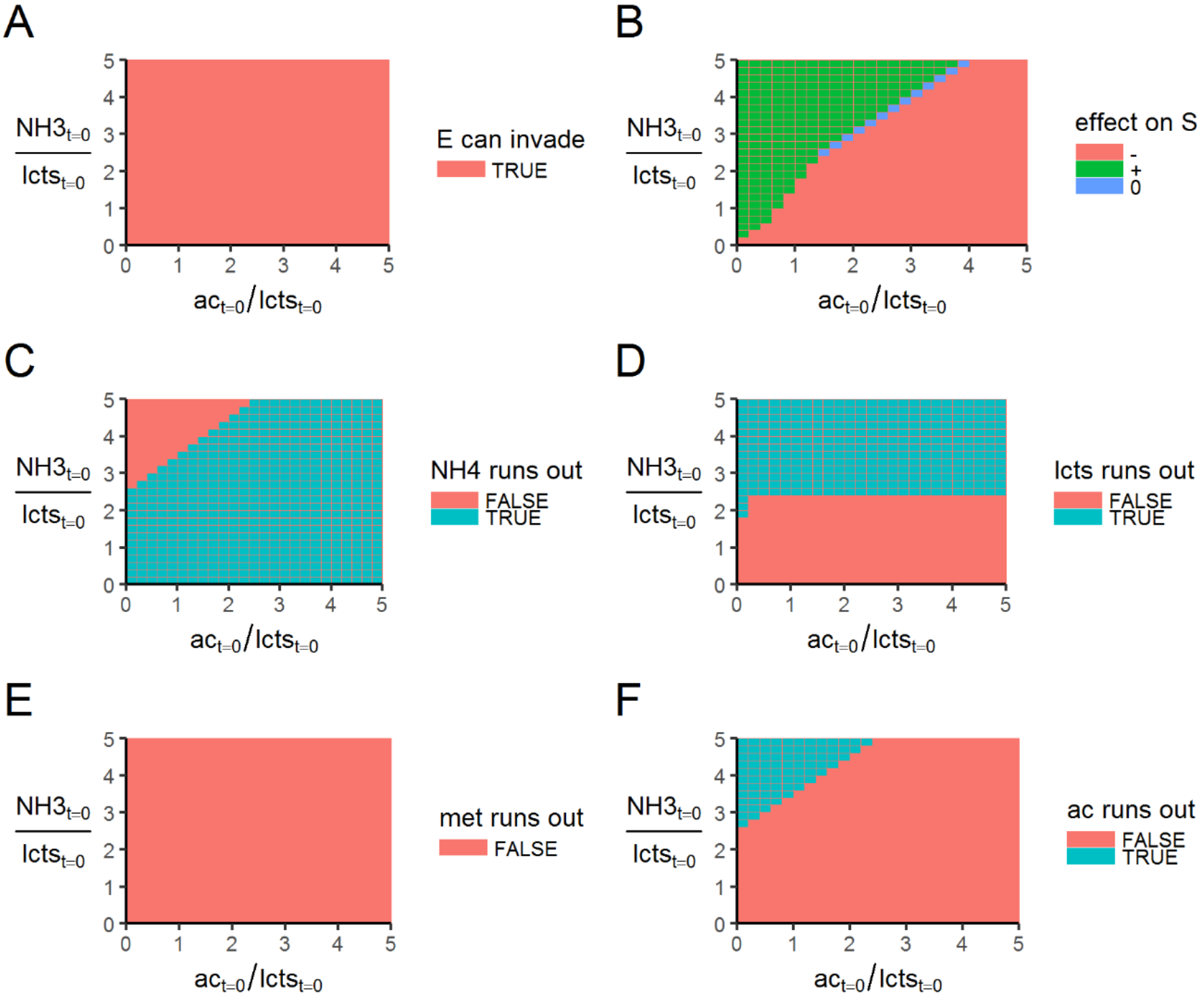
Simulated effect of initial acetate and NH_3_ concentrations on the ability of a faster-growing obligate cross-feeding species to coexist with its nutrient supplier. Results of simulations across many initial values of NH_3_ and ac. In all simulations, *S*_t=0_ = 198, *E*_t=0_ = 2, lcts_t=0_ = 10000, μ_Smax_ = 0.5, and λ_ac_ = λ_met_ = 1.5. **(A)** Whether *E* can increase in frequency and invade from rare. **(B)** The net effect of *E*’s presence on *S*, calculated by comparing *S*’s yield in the presence / absence of *E*. The net effect of *S* on *E* due to *E*’s dependence on met, which is absent in the environment. **(C-F)** show whether different nutrients run out in the simulations (**(C)** = NH_3_, **(D)** = lcts, **(E)** = met, **(F)** = ac).

## Supplemental Experimental Procedures

### Scaling of the General Ecological Model

#### 1. The original model

First, the growth rates functions, which specify biomass growth of each species as a function of that species’ population size and the concentrations of their required nutrients.

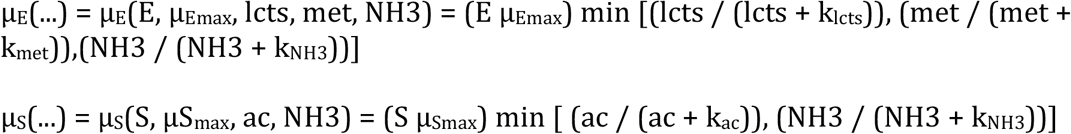

In the above equations, the k_x_ parameters signify the concentration of x at which growth is half-maximum. The μ_Xmax_ parameters signify the maximum possible growth rate (/hr) species x can attain.

The above growth rate functions drive each species’ growth:

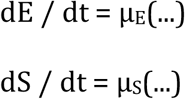

Resources are either consumed during growth or produced during growth:

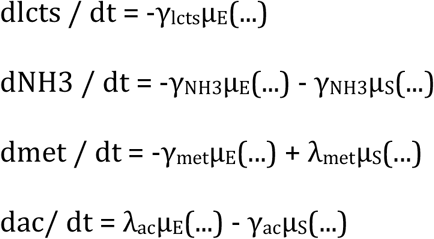

In the resource equations, the γ_x_ parameters signify the number of units of resource x which are *consumed* during growth of one unit of the relevant species, and the λ_x_ parameters signify the number of units of resource x which are *produced* during growth of one unit of the relevant species.

The species E and S are in units of cells, the resources are in units of grams, and therefore the γs and λs are in units of grams per cell.

#### 2. The scaling

First, we scale the resources from units of grams / cell, to equivalent cell units, by dividng the resources by the associated consumption parameter γ:

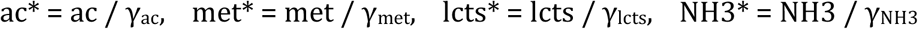

Next, we scale the production parameters (λs) by the consumption parameters (γs), which allows us to consider production in terms of the number of cells of one species which can be supported by the growth of one cell of the other species:

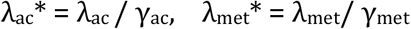

These scaled production parameters are in units of cells / cell, i.e. are unitless. For intuition, if λ_ac_* = 2, then for each cell of E grown, enough acetate is produced to support growth of 2 S cells.

Next, the half-saturation parameters (k) must be scaled to keep the units consistent. This is also done by scaling by the consumption parameters (γs), such that the scaled half-saturation parameters describe how many cells’ worth of resources are present when growth is at half-maximum:

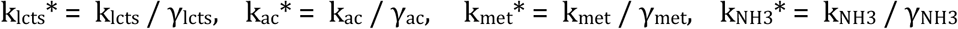

Using the chain rule and substitutions of the above scaled parameters, we can remove the consumption terms from our resource equations, for example resulting in the ammonia equation now specified as:

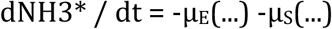

The growth rate functions (μ_x_(…)) were similarly adjusted, to take in scaled resource variables and half-saturation parameters.

The final scaling we do is to scale time by E’s maximum growth rate. We do this because we are interested in the effect of relative growth rates between E and S, not their absolute growth rates, which only change the time scale but not the species ratios. We scale time by doing:

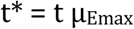

And then scale S’s maximum growth rate by E’s:

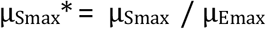

With these scalings we can remove E’s maximum growth rate and are only left with the relative growth rate μ_Smax_* in S’s growth equation. For clarity, in the main text, we omit the asterisks from variable names.

